# Ethanol-induced conditioned taste aversion and associated neural activation in male rats: Impact of age and adolescent intermittent ethanol exposure

**DOI:** 10.1101/2022.05.31.494165

**Authors:** Jonathan K. Gore-Langton, Elena I. Varlinskaya, David F. Werner

**Author notes:** Address Correspondence to David F. Werner, Ph.D., Department of Psychology, Binghamton University-State University of New York, Binghamton, NY 13902-6000.

## Abstract

Individuals that initiate alcohol use at younger ages and binge drink during adolescence are more susceptible to developing alcohol use disorder. Adolescents are relatively insensitive to the aversive effects of alcohol and tend to consume significantly more alcohol per occasion than adults, an effect that is conserved in rodent models. Adolescent typical insensitivity to the aversive effects of alcohol may promote greater alcohol intake by attenuating internal cues that curb its consumption. Attenuated sensitivity to the aversive effects of alcohol is also retained into adulthood following protracted abstinence from adolescent intermittent ethanol (AIE) exposure. Despite these effects, much remains unknown regarding the neural contributors. In the present study, we used a conditioned taste aversion (CTA) paradigm to investigate neuronal activation in late-developing forebrain structures of male adolescents and adult cFos-LacZ transgenic rats as well as in AIE adults following consumption of 0.9% sodium chloride previously paired with an intraperitoneal injection of 0, 1.5 or 2.5g/kg of ethanol.. Unlike adults that were non-manipulated or received water exposure during adolescence, adolescents as well as adults who experienced AIE did not display CTA to a 1.5 g/kg ethanol dose. Adults displayed increased neuronal activation indexed via number of β-galactosidase positive (β-gal+) cells in the prefrontal and insular cortex that was absent in adolescents, whereas adolescents but not adults had reduced number of β-gal+ cells in the central amygdala. Adults also displayed greater cortical-insular functional connectivity than adolescents as well as insular-amygdalar and prefrontal cortex-accumbens core functional connectivity. Like adolescents, adults previously exposed to AIE displayed reduced prefrontal-insular cortex and prefrontal-accumbal core functional connectivity. Taken together, these results suggest that attenuated sensitivity to the aversive effects of ethanol is related to a loss of an insular-prefrontal cortex-accumbens core circuit.

## Introduction

Alcohol is the most used drug among adolescents worldwide. According to Monitoring the Future survey, in 2020 more than half (55.3%) of high school seniors had used alcohol in the past year and 16.8% drank at binge levels within the past two weeks (Johnston et al., 2021). Rates of past-year alcohol use and binge drinking, defined by the National Institute of Alcohol Abuse and Alcoholism as a pattern of drinking that brings blood alcohol concentrations to 0.08 g/dl or above, were lower for 10^th^ graders (40.7 and 9.6%) and 8^th^ graders (20.5 and 4.5%). Several studies have shown that early onset of alcohol use as well as an escalation of drinking during adolescence increases the risk of developing an alcohol use disorder (AUD) in adulthood (Elsayed et al., 2018; Sartor et al., 2016; Waller et al., 2019). According to the Substance Abuse and Mental Health Services Administration (Bose, 2016), adolescents and adults display different patterns of alcohol drinking: while adults consume alcohol more frequently than adolescents, adolescents tend to drink substantially more per occasion, with some adolescents demonstrating high-intensity or extreme binge drinking by consuming 10+ and even 15+drinks in a row (Patrick & Terry-McElrath, 2019; Patrick et al., 2013). It is unfortunate that alcohol use is relatively high among adolescents, given that this demographic is particularly susceptible to the long-term negative neurocognitive and neurodevelopmental effects of alcohol. For example, binge drinking during adolescence has been associated with impairment of attention, information retrieval, and visuospatial skills (Brown et al., 2000). Magnetic resonance imaging (MRI) studies have shown that hippocampal volumes tend to be smaller in adolescents with AUDs than age-matched controls and that this effect is more pronounced with earlier alcohol onset and greater duration of drinking (De Bellis et al., 2000; Welch et al., 2013).

Given the established harms of adolescent alcohol use, it is important to characterize neural contributors to adolescent-typical binge and high-intensity drinking and to investigate alcohol-induced alterations in the developmental trajectory of the adolescent brain. Therefore, animal models (typically rodents) are a useful tool for understanding adolescent-typical responsiveness to ethanol as well as the neural perturbations caused by adolescent ethanol exposure, which is often not possible with human subjects. Animal models allow for control of a number of variables, including genetic background, environmental conditions, and ethanol exposure regimens (dose, frequency, age, and duration of exposure), while manipulating only variables of interest (e.g., age, sex, early experience, etc). Animal studies recapitulate that adolescent rats and mice not only consume more ethanol on a g/kg basis than their adult counterparts (Doremus et al., 2005; Salguero et al., 2020; Spear, 2014; Varlinskaya et al., 2015; Vetter-O’Hagen et al.,2009), but also are less sensitive than adults to the undesired effects of ethanol that may serve as cues to curb drinking. These adverse effects include ethanol-induced motor impairment (Ramirez & Spear, 2010; White, Bae et al., 2002; White, Truesdale et al., 2002), sedation (Draski et al., 2001; Silveri & Spear, 1998), social inhibition (Varlinskaya & Spear, 2002), and aversion (Anderson et al., 2010; Saalfield & Spear, 2016; Schramm-Sapyta et al., 2014; Vetter-O’Hagen et al., 2009). In laboratory rodents, sensitivity to the aversive effects of ethanol has commonly been assessed using a conditioned taste aversion (CTA) paradigm. In this paradigm, ingestion of a novel flavour (conditioned stimulus; CS) is paired with the dysphoric effects of ethanol or other drugs (unconditioned stimulus; US). When an animal is given a subsequent opportunity to consume the CS, it will tend to avoid or reduce its intake, presumably due to the association of the CS with the aversive effects of the drug. The degree to which the animal reduces its consumption relative to a vehicle-injected control is used as an index of the dysphoria experienced during the initial CS-US pairing. Relative to adults, adolescent rats require higher doses (Anderson et al., 2010; Saalfield & Spear, 2016) or more CS-US pairings to develop a CTA (Anderson et al., 2010). As the overall hedonic value of a drug is thought to be a function of the balance between its rewarding and aversive effects (Davis & Riley, 2010; Riley, 2011), the relative insensitivity of adolescents to the aversive effects of ethanol may contribute to high levels of adolescent ethanol intake. Indeed, a meta-analysis of rodent genetic studies confirmed that insensitivity to the aversive effects of ethanol assessed in a CTA paradigm was more strongly related to EtOH consumption than sensitivity to the rewarding effects (Green & Grahame, 2008).

Similar insensitivity to the adverse effects of ethanol has been reported in adult laboratory rodents following chronic exposure to ethanol during adolescence. Adult mice and rats exposed to ethanol in adolescence demonstrated attenuated sensitivity to ethanol-induced sedation (Matthews et al., 2008), motor impairment (White, Bae et al., 2002), and aversion (Diaz-Granados & Graham, 2007; Saalfield & Spear, 2015). It is likely that attenuated sensitivity to some adverse effects of ethanol is rather specific to adolescent ethanol exposure, since attenuation of ethanol sensitivity was not evident following equivalent ethanol exposure and abstinence in adulthood (Diaz-Granados & Graham, 2007; White et al., 2000). Furthermore, attenuated sensitivity to ethanol is more pronounced when adolescent ethanol exposure is intermittent rather than continuous, mimicking adolescent-typical binge drinking (see Spear, 2020 for review). Therefore, adolescent intermittent ethanol (AIE) exposure of laboratory rodents is frequently used as an animal model of adolescent binge-like episodes (Crews et al., 2019; Marco et al., 2017). Route of ethanol administration does not play a substantial role, given that attenuated sensitivity to different ethanol effects in adulthood has been reported following ethanol vapor inhalation (Diaz-Granados & Graham, 2007), intraperitoneal injection (Matthews et al., 2008), and intragastric gavage (Saalfield & Spear, 2015).

Understanding the neural contributors of adolescent typical resistance to the aversive effects of ethanol as well as AIE-associated insensitivity to ethanol-induced CTA may prove important for understanding mechanisms of high-intensity drinking during adolescence and the development of AUDs in later life. To our knowledge, only one study has assessed patterns of neuronal activation using Fos-immunoreactivity in adolescent and adult males following ethanol-induced CTA in response to the US (Saalfield & Spear, 2019). Although AIE has been shown to attenuate ethanol-induced CTA in male mice (Diaz-Granados & Graham, 2007) and rats (Saalfield & Spear, 2015), no studies to our knowledge have investigated patterns of neuronal activation in CTA-associated brain regions in adults following AIE exposure. Therefore, the present experiments were designed to assess patterns of neuronal activation within brain regions involved in response to the CS following ethanol-induced CTA in adolescent and adult male rats (Experiment 1) as well as in adult males following AIE (Experiment 2).

Male cFos-LacZ transgenic rats on a Sprague-Dawley background were used as experimental subjects. Since these transgenic rats express β-galactosidase (β-gal) under the control of the *cfos* promoter, cFos and β-gal proteins are co-expressed in neurons strongly activated by a certain stimulus and not in the surrounding neurons that are inactive or weakly activated (Cruz et al., 2014). Therefore, β-gal expression can be used as a proxy for cFos. While cFos expression requires immunohistochemical techniques, β-gal is detected by X-gal staining – a rapid and convenient histochemical assay. In Experiment 1, the regions of interest (ROIs) included the prelimbic cortex (PrL), infralimbic cortex (IL), agranular insula ventral (AIV) and dorsal (AID) subdivisions, nucleus accumbens core (NAcC) and shell (NAcSh), basolateral (BLA) and central (CeA) amygdala. We focused on later developing cortical and subcortical regions (e.g., PFC, NAcC) involved in CTA learning since we hypothesized that these would be more likely to show age-related differences in neuronal activity compared to earlier developing brainstem and more caudal structures (e.g., NTS, PBN). The dorsal striatum (DS) which has no known role in CTA was included as a control ROI. In Experiment 2, neuronal activation by the CS was assessed in the AIV, AID, PrL, IL, NAcC, BLA, and CeA.

## General Methods

### Subjects

All experimental subjects were generated from cFos-LacZ transgenic male rats that were backcrossed with outbred Sprague-Dawley females from our animal colony at Binghamton University over multiple generations. Breeders were obtained from the transgenic line of animals originally developed by Dr. Curran while at Roche Institute of Molecular Biology. Experimental animals were from the N7-8 generation (99.9% gene similarity) with our standard Sprague-Dawley breeders or higher. One day after birth [postnatal day (P)1], litters were culled to 8-10 pups, with no more than a 2-pup difference between sexes. On P21, pups were weaned and group-housed with same sex littermates. Between P20 and P24 tissue was collected by ear punch for LacZ genotyping (TransnetYX, Cordova, TN). Only LacZ positive (LacZ+) males were used. Animals were housed in a temperature-controlled (22°C) vivarium on a 12/12 light/dark cycle (lights on at 0700). All animals were treated under guidelines established by the National Institutes of Health and protocols approved by Binghamton University Institutional Animal Care and Use Committee.

### Conditioned Taste Aversion (CTA)

The day before conditioning, animals were single-housed and given 50% of the water they consumed the previous day to encourage intake of the CS on conditioning day. On conditioning day, animals were transported in their home cages from the holding room to a testing room containing dim light (15-20 lux) and a white noise generator set at 68 decibels. Home cage tops containing food were replaced with new tops without food. After 15 min of habituation to the room, a single bottle containing 0.9% sodium chloride (NaCl) was placed on the home cage for a 1-h access period. Then bottles were removed and weighed, and animals received an i.p. injection of a single ethanol dose. Animals that drank *less than 0*.*5 g of NaCl* were excluded. Following the EtOH injection, animals were left in the testing room for another 15 min prior to being returned to their holding rooms. At this time, home cage tops were returned, and animals were given full water bottles. The next day animals were given 50% of the water they consumed in their home cage within the past 24 h. The following day CTA testing occurred, with animals again being exposed to the CS (NaCl) for 1 h. After removal of the bottles, rats were left undisturbed for 1 h until they were euthanized for tissue collection.

### Tissue Preparation

Animals were euthanized with sodium pentobarbital (Fatal-Plus, Vortech Pharmaceuticals, Dearborn, MI) and perfused with 0.1M phosphate-buffered saline (PBS) at a rate of 30ml/min (200cc for adolescents or 250cc for adults). Then the flow was switched to 4% paraformaldehyde in 0.1M PBS (infused at same volume as PBS) to fix the tissue. Immediately after fixation, the brain was extracted and placed into a vial containing 4% paraformaldehyde solution for a 90 min post-fix period. The brain was then transferred into a cryoprotective solution of 30% sucrose in water until saturated, and temporary storage at −20°C until the brain was sectioned and stained.

All collected brains prior to being coronally sectioned were flash-frozen in methyl-butane at −80°C. Once brains were prepared to be sectioned the cerebellum was removed with a razor blade and the brain was mounted to a chuck by applying OCT gel and frozen between temperatures of −20 to −16°C prior to slicing. Slices were collected every 30µm and transferred into 1.7ml microcentrifuge tubes containing antifreeze solution (30% ethylene glycol and 30% glycerol in 0.5M PBS) then stored at −20°C until X-Gal histochemistry was performed. A serial collection method was used in which every 12^th^ slice (360µm between each collection) was placed into the same tube for a total of 6 slices per tube.

### X-Gal Histochemistry

On the first day of the X-Gal enzymatic reaction procedure, representative brain regions were gently agitated in X-Gal fix buffer (2mM magnesium chloride and 5mM EGTA in 0.1M PBS) followed by two washes in X-Gal wash buffer (2mM magnesium chloride in 0.1M PBS). Next, slices were placed into X-Gal staining solution (2mM magnesium chloride, 5mM potassium ferrocyanide, 5mM potassium ferricyanide, and 1 mg/ml of X-Gal in 0.1M PBS) and left to incubate overnight in a 37°C orbital stage chamber. On the second day, slices were again washed twice in X-Gal wash buffer and mounted onto chromalum-gelatin-coated slides and left to air-dry over night at room temperature. The next day, mounted slices were counterstained in eosin and dehydrated using a series of ascending percentages of, then cleared with Xylene and coverslipped.

### Regions of Interest (ROI) and Cell Counting

In Experiment 1, imaging was conducted using a Keyence BZ-X800 microscope (Keyence corporation, Itasca, IL) and stained soma were quantified using NIH Image J software. ROIs included PrL, IL, AIV, AID, NAcC, NAcSh, BLA, CeA, and DS. β-gal+ cells were manually counted by the same experimenter within a set size grid box for each ROI (see Table 1). The rat brain atlas (Paxinos and Watson, 2007) was used as a guide for grid placement and size. All cells found on the border of the grid box (i.e., partially obscured by border) were counted. In Experiment 2, ROIs included the PrL, IL, AIV, AID, NAcC, BLA, and CeA. Imaging was done using an Olympus VS200 slide scanner (Olympus corporation, Tokyo, Japan) and stained soma were quantified using HALO 3.0 cytonuclear module (Indica Labs, Albuquerque, NM). Parameters for the HALO analysis algorithm for detection of a β-gal+ expressing cell were set at a minimum optical tissue density of 0.037 and minimum nuclear area of 10. Comparisons between cell counts between manual counting methods versus Halo did not reveal any significant differences in any of the assessed regions. All β-gal+ cell counts were standardized to an area of 1mm^2^ to make graphical comparisons between cell density in different ROIs

**Table 1.** Brain Region Coordinates Based on the Paxinos and Watson Atlas (Paxinos & Watson, 2007)

| ROI | Grid Size | Coordinates from Bregma |
| --- | --- | --- |
| AIV | 0.2 X 1mm (0.2mm<sup>2</sup>) | 4.20 to 2.28mm |
| AID | 0.2 X 1mm (0.2mm<sup>2</sup>) | 4.20 to 2.28mm |
| PrL | 0.5mm<sup>2</sup> | 5.16 to 2.76mm |
| IL | 0.25mm<sup>2</sup> | 3.72 to 2.76mm |
| NAcC | 0.4mm<sup>2</sup> | 2.76 to 0.72mm |
| NAcSh | 0.4mm<sup>2</sup> | 2.52 to 0.72mm |
| BLA | 0.5mm<sup>2</sup> | -1.72 to -3.24mm |
| CeA | 0.5mm<sup>2</sup> | -1.72 to -3.00mm |
| DS | 0.5mm<sup>2</sup> | 2.52 to 0.00mm |

### Experiment 1: Design and Statistical Analyses

Experiment 1 was designed to assess the impact of age on neuronal activation in CTA-associated brain regions following re-exposure to CS. The design was a 2 Age (adolescent, adult) X 3 Ethanol Dose (0,1.5, 2.5) factorial. Animals were single-housed on P25, for animals tested in adolescence and P65 for animals tested in adulthood. Conditioning occurred on P30 (adolescents) or P70 (adults), with CTA testing and brain tissue collection occurring two days later on P32 or P72 (see Figure 1a).

**Figure 1.**
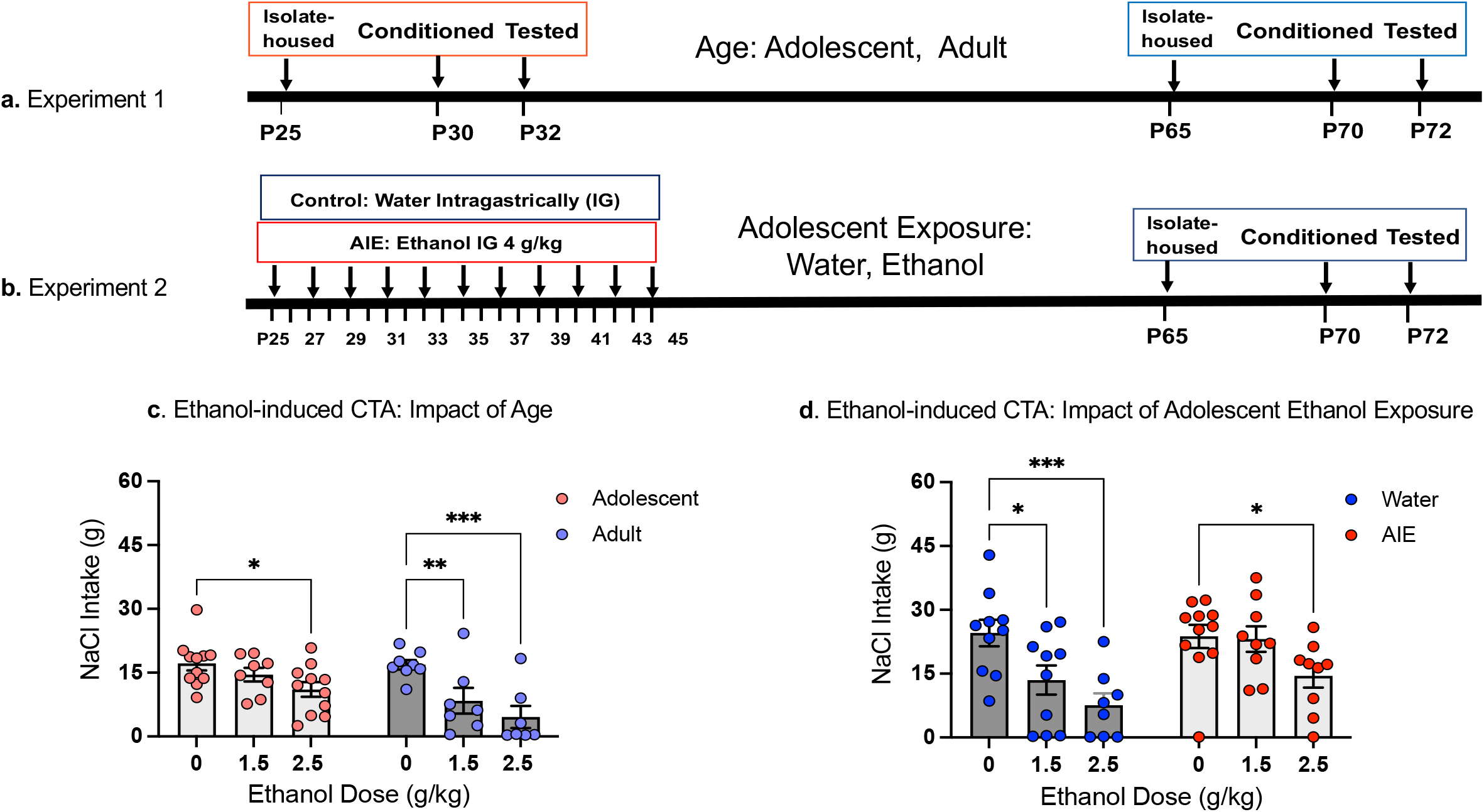
Timelines and NaCl (CS) intake on test day in Experiment 1 (a, c) and Experiment 2 (b, d). Asterisks depict significant changes in NaCl intake on test day within each Age relative to controls conditioned with saline (0 g/kg EtOH dose), * - p < 0.05, ** - p <0.01, *** - p < 0.001.

Intake of the CS on both conditioning and test day were recorded in grams (g) of fluid and assessed using 2 Age (i.e., adolescent, adult) X 3 Ethanol Dose (0,1.5, 2.5 g/kg) ANOVAs, with Fisher’s pair-wised post-hoc tests used to determine ethanol doses that induced CTA. Development of a CTA was determined as a significant decrease in intake from the age-appropriate saline (0 g/kg) control group. β-gal expression was examined in each ROI using a 2 Age X 3 Ethanol Dose factorial ANOVA. In the case of age differences indexed via a main effect of Age, omnibus ANOVAs were followed by separate one-way ANOVAs for each age. To explore the locus of main effects or their interactions, Fisher’s LSD tests were used. A forward multiple stepwise regression was conducted separately for each age. allowing for investigation of the influence of each brain region (i.e., β-gal expression) on the strength of CTA (i.e., test day intake). Patterns of neuronal activation across brain regions were investigated using Pearson bivariate correlations between β-gal in each region within each Age. A false discovery rate of 5% (q-value threshold = 0.05) was maintained using the two-stage set up method of Benjamini, Krieger and Yekutieli to limit the number of false positive comparisons. One outlier animal (adolescent 0 g/kg animal that was +2 SDs greater than the group mean in several distinct ROIs (AID, AIV, IL, NAcSh, BLA) was excluded from the correlation matrix analysis.

All analyses were separated by Age as factorial ANOVAs revealed greater β-gal expression in adolescents compared to adults in all ROIs except for the BLA where there was a trend for greater expression in adolescents (*p*=0.08).

### Experiment 2: Design and Statistical Analyses

Experiment 2 was designed to assess the effect of adolescent intermittent ethanol (AIE) exposure on neuronal activation in CTA-associated brain regions following re-exposure to the CS. The design was a 2 Adolescent Exposure (water, AIE) X 3 Ethanol Dose (0,1.5, 2.5) factorial. Experimental subjects were given either tap water or 4 g/kg ethanol (AIE) intragastrically (25% v/v in tap water) every other day between P25 and P45 for a total of 11 exposures. All animals that were housed in the same cage received the same exposure (water or AIE). After the last exposure on P45, experimental subjects were left undisturbed for 20 days. Animals remained group-housed with no more than 4 animals per cage until P65 at which point, they were single-housed. Animals were conditioned on P70 and tested for CTA on P72 (see Figure 1b).

Body weights of Water and AIE exposure groups were compared across Days (P25-45) using a repeated measures ANOVA. Intake of the CS on both conditioning and test day were assessed by separate 2 Adolescent Exposure (Water, AIE) X 3 Ethanol Dose (0,1.5, 2.5 g/kg) ANOVAs. Fisher’s LSD tests were used to determine effective doses of ethanol within each adolescent condition for producing CTA. Number of β-gal+ cells was examined in each ROI using a 2 Adolescent Exposure (Water, AIE) X 3 Ethanol Dose (0,1.5, 2.5 g/kg) ANOVA. Omnibus ANOVAs were followed by separate one-way ANOVAs for each Exposure in the case of main effects of Exposure and/or EtOH Dose. In the case of main effects of Adolescent Exposure and/or Ethanol Dose, separate one-way ANOVAs for each Exposure were used followed by Fisher’s LSD *post-hoc* tests. A forward multiple stepwise regression was conducted separately for each Adolescent Exposure condition in order to assess the influence of each brain region (i.e., β-gal expression) on the strength of CTA (i.e., test day intake). All results were considered statistically significant at *p*≤0.05. Finally, potential functional connectivity between brain regions was investigated using Pearson bivariate correlations between β-gal in each region for each Exposure. A false discovery rate of 5% (q-value threshold = 0.05) was maintained using the two-stage set up method of Benjamini, Krieger and Yekutieli to limit the number of false positive comparisons.

## Results

### Experiment 1. Conditioned taste aversion and neuronal activation in CTA-associated brain regions following re-exposure to CS: Impact of age

#### Ethanol-induced CTA

The intake of sodium chloride (CS) on conditioning day did not differ as a function of Age and Ethanol Dose (all p >0.05), with adolescents and adults ingesting comparable amounts of the CS (14.4 ± 0.7 g and 16.1± 1.6 g, respectively). On test day, the 2 Age X 3 Ethanol Dose ANOVA of the CS intake revealed main effects of Age (*F*_1,46_ = 7.12, *p*< 0.05) and Ethanol Dose (*F*_2,46_ = 11.75, *p*<0.01), with adolescents consuming significantly more of the CS on test day (14.23 ± 1.06 g) than adults (10.26 ± 1.70 g) when data was collapsed across Ethanol Dose (*t*_50_ = 2.08, *p* < 0.05). Follow-up separate for each age one-way ANOVAs revealed a main effect of Ethanol Dose for both adolescents (*F*_2,27_ =3.67, *p*<0.05) and adults (*F*_2,19_ =7.71, *p*<0.01). Post-hoc tests showed that adults developed CTA to both the 1.5 and 2.5 g/kg ethanol doses, demonstrating significantly lower intake of NaCl than saline control group, while in adolescents, CTA was evident only following the 2.5 g/kg dose (Figure 1c). These results indicate that adolescent cFos-LacZ transgenic male rats were less sensitive to ethanol-induced CTA than their adult counterparts.

#### Neuronal Activation

##### Prelimbic Cortex (PrL) and Infralimbic Cortex (IL)

The ANOVA of β-gal expression in the PrL revealed a main effect of Age (*F*_1,46_ = 5.09, *p*<0.01), with significantly more β-gal+ cells evident in the PrL of adolescents than adults (218.04 ± 12.75 and 180.31 ± 10.87, respectively, *t*_50_ =2.15, *p* < 0.05, see Figure 2a). Follow-up one-way ANOVAs revealed a main effect of Ethanol Dose in adults (*F*_2,19_ =3.96, *p*<0.05), but not adolescents. Adults that received the 1.5 g/kg EtOH dose on conditioning day had significantly more β-gal+ cells in the PrL than saline controls (see Figure 2a).

**Figure 2.**
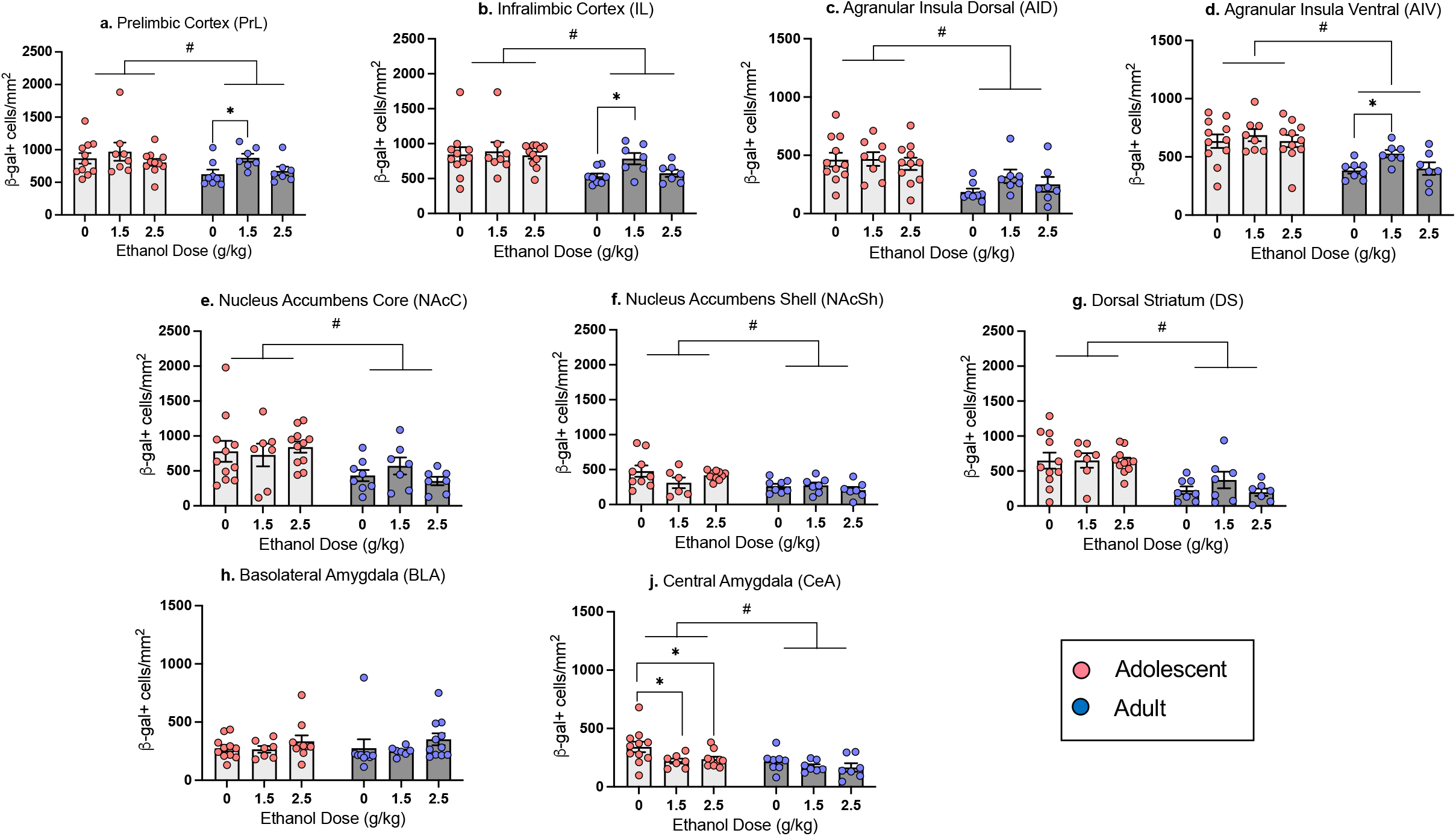
Neuronal activation indexed via number of β-gal+ cells in the Prelimbic Cortex (a), Infralimbic Cortex (b), Dorsal Agranular Insular Cortex (c), Ventral Agranular Insular Cortex (d), Nucleus Accumbens Core (e), Nucleus Accumbens Shell (f), Dorsal Striatum (g), Basolateral Amygdala (h), and Central Amygdala (j) in adolescent and adult males previously conditioned with 0, 1.5 or 2.5 g/kg ethanol on test day following exposure to the CS (NaCl). n = 7-11/group; # denotes main effect of age, asterisks (*) mark significant ethanol-associated changes relative to controls conditioned with with saline (0 g/kg ethanol dose) within each Age (p < 0.05).

In the IL, the omnibus ANOVA revealed a main effect of Age (*F*_1,46_ =9.95, *p*<0.01), with a greater number of β-gal+ cells in adolescents than adults (52.16 ± 3.05 and 39.14 ± 2.50, respectively, *t*_50_ =3.14, *p* < 0.01, Figure 2b). One-way ANOVAs revealed a main effect of Ethanol Dose in adults (*F* _2,19_ =5.29, *p*<0.050, but not adolescents, with adults in the 1.5 g/kg dose group demonstrating significantly more β-gal+ cells relative to the saline group, with no changes evident in the 2.5 g/kg ethanol dose condition (see Figure 2b).

##### Dorsal (AID) and Ventral (AIV) Agranular Insular Cortex

In the AID, the two-factor ANOVA revealed a main effect of Age (*F*_1,46_ =18.62, *p*<0.01), with a significantly higher number of β-gal+ cells in adolescents than adults (90.22 ± 6.42 and 49.84 ± 6.07, respectively, *t*_50_ = 4.42, *p* < 0.001, Figure 2c). One-way ANOVAs did not reveal an effect of Ethanol Dose in either adolescents or adults. Similarly, the factorial ANOVA of β-gal expression in the AIV revealed a significant main effect of Age (*F*_1,46_ = 26.13, *p*<0.01). The number of β-gal+ cells in the adolescent AIV was significantly higher than in the adult AIV (129.87 ± 6.22 and 86.91 ± 5.04, respectively, *t*_50_ =5.08, *p* < 0.001, Figure 2d). One-way ANOVAs within each Age revealed a main effect of Dose only in adults (*F* _2,19_ =4.21, *p*<0.05), with adults conditioned with the 1.5 g/kg ethanol dose demonstrating significantly more β-gal+ cells than saline controls.

##### Nucleus Accumbens Core (NAcC), Nucleus Accumbens Shell (NAcSh), and Dorsal Striatum (DS)

In the NAcC, the factorial ANOVA revealed a main effect of Age (*F*_1,45_ =11.02, *p*<0.01), with β-gal+ cell counts in adolescents being significantly higher than in adults (126.57 ± 11.70 and 72.45 ± 8.52, respectively, *t*_49_ =3.52, *p* < 0.001, Figure 2e). One-way ANOVAs did not reveal effects of Ethanol Dose at either age. In the NAcSh, there was a main effect of Age (*F*_1,42_ = 13.04, *p*<0.01) as well, with adolescents demonstrating significantly higher β-gal+ cell counts than adults (68.35 ± 5.60 and 39.76 ± 3.62, respectively, *t*_46_ = 3.92, *p* < 0.001, Figure 2f). Follow-up ANOVAs did not reveal effects of Ethanol Dose in either adolescents or adults. In the DS, β-gal+ cell counts only differed as a function of Age (*F* _1,45_ =25.93, *p*<0.01), with adolescents demonstrating higher β-gal expression than adults (161.40 ± 13.05 and 65.79 ± 11.63, respectively, *t*_49_ =5.28, *p* < 0.001, Figure 2g). One-way ANOVAs did not reveal effects of Ethanol Dose in either adolescents or adults.

##### Basolateral Amygdala (BLA) and Central Amygdala (CeA)

In the BLA, β-gal+ cell counts did not differ as a function of Adolescent Exposure or Ethanol Dose (see Figure 2h). In the CeA, the ANOVA of β-gal expression revealed a main effect of Age (*F*_1,44_ =8.96, *p*<0.01) and Ethanol Dose (*F*_2,44_ =3.75, *p*<0.05). One-way ANOVAs showed a main effect of Dose (*F*_2,25_ =3.45, *p*<0.05) in adolescents, but not in adults, with adolescents previously conditioned with either the 1.5 or 2.5 g/kg ethanol showing significant decreases in β-gal+ cells relative to controls conditioned with saline (Figure 2j).

##### Forward-Stepwise Multiple Regression: β-gal and Test Day intake

In adolescents the forward stepwise multiple regression model revealed a significant positive relationship between β-gal expression in the AID and test day intake (adjusted R^2^=0.136, *p*=0.028). In other words, greater β-gal expression in the AID of adolescents was related to greater intake of the CS on test day. There were no other significant predictors of test intake in the adolescent model. In adults, there was a significant negative relationship between β-gal expression in the BLA and test day intake (adjusted R^2^=.134, *p*=0.025). All other ROIs did not differ from the model by the stepwise procedure.

##### Patterns of neuronal activation: Correlation matrix

Pearson bivariate correlations were conducted using pairwise comparisons of β-gal+ cell counts in each ROI (see Figure 4a). In both adolescents and adults, significant positive correlations (all *p* < 0.05) were evident between the PrL and IL, PrL and AIV, IL and AIV, AID and AIV, as well as NAcC and NAcSh. Adults displayed increased functional connectivity over adolescents, with significant positive correlations evident between the IL and AID, IL and NAcC, IL and DS, AID and BLA, AIV and BLA, NAcC and DS, NAcSh and DS.

### Experiment 2. Conditioned taste aversion and neuronal activation in CTA-associated brain regions following re-exposure to CS: Impact of prior history of adolescent intermittent ethanol (AIE) exposure

#### Ethanol-induced CTA

The intake of sodium chloride (CS) on conditioning day differed as a function of Adolescent Exposure (F_1,51_ = 6.18, *p* < 0.05), with AIE-exposed males consuming significantly (*t* _55_ = 2.41, *p* < 0.05) more than their water-exposed counterparts (25.43g ± 1.35g and 20.39g ± 1.60g, respectively). On test day, the 2 Adolescent Exposure X 3 Ethanol Dose ANOVA of CS intake revealed significant main effects of Adolescent Exposure (*F*_1,51_ = 4.60, *p*<0.05) and Ethanol Dose (*F* _2,51_ = 9.38, *p*<0.01), with AIE animals consuming more than water-exposed controls and rats that were conditioned either with the 1.5 or 2.5 g/kg dose demonstrating significantly lower intake of the CS at test relative to those conditioned with the 0 g/kg ethanol dose. Follow-up one-way ANOVAs conducted within each Adolescent Exposure revealed a main effect of Ethanol Dose for both AIE-exposed (*F*_2,26_ =3.26, *p*=0.054) and water-exposed (*F* _2,25_ =7.15, *p*<0.01) animals. P*ost-hoc* tests showed that AIE-exposed rats only developed CTA at the 2.5 g/kg EtOH dose, whereas water-exposed controls demonstrated CTA at both the 1.5 and 2.5 g/kg EtOH (see Figure 1d), suggesting that AIE exposure attenuated sensitivity to EtOH-induced CTA in adulthood.

#### Neuronal Activation

##### Prelimbic Cortex (PrL) and Infralimbic Cortex (IL)

The number of β-gal+ cells in the PrL did not differ as a function of Adolescent Exposure and Ethanol dose (see Figure 3a). However, in the IL a two-way ANOVA revealed a main effect of Adolescent Exposure (*F*_1,50_ = 8.21, *p* < 0.01), with AIE-exposed animals demonstrating more β-gal+ cells than their water-exposed counterparts (60.22 ± 2.31 and 51.89 ± 1.73, respectively), and a main effect of Ethanol Dose (F_2,50_ =4.99, *p* <0.05], with significant decreases in β-gal expression evident in animals conditioned with either the 1.5 or 2.5 g/kg EtOH dose when collapsed across Adolescent Exposure. Follow-up one-way ANOVAs within each Exposure group did not reveal effects of Dose in the water exposed group and only a suggestive effect in the AIE exposed group (*p*=0.057) (See Figure 3b).

**Figure 3.**
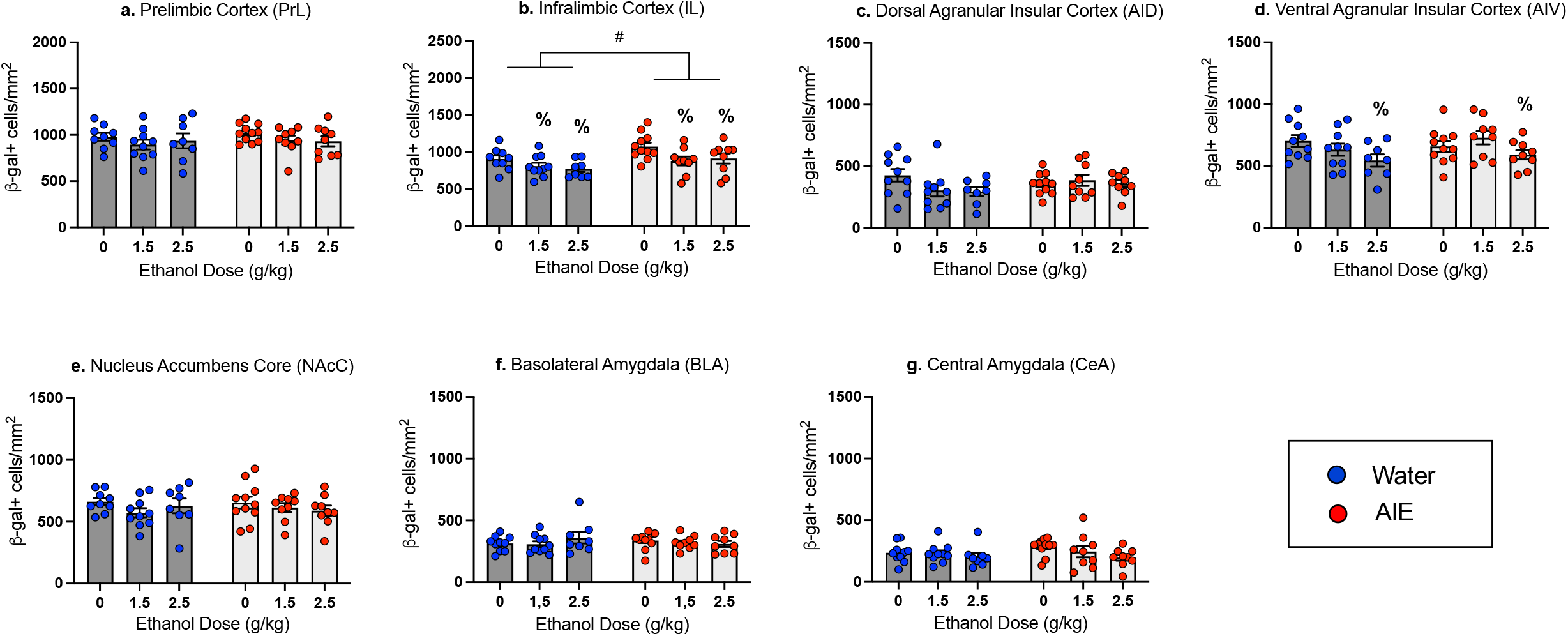
Neuronal activation indexed via number of β-gal+ cells in the Prelimbic Cortex (a), Infralimbic Cortex (b), Dorsal Agranular Insular Cortex (c), Ventral Agranular Insular Cortex (d), Nucleus Accumbens Core (e), Basolateral Amygdala (f), and Central Amygdala (g) in adult males exposed to water or ethanol (AIE) during adolescence and previously conditioned with 0, 1.5 or 2.5 g/kg ethanol on test day following exposure to the CS (NaCl). n = 8-10/group; average number of slices per region for individual animal = 3-7. # denotes main effect of Adolescent Exposure, %marks significant ethanol-associated changes relative to controls conditioned with with saline (0 g/kg ethanol dose) when collapsed across Adolescent Exposure (p < 0.05).

##### Dorsal (AID) and Ventral (AIV) Agranular Insular Cortex

The factorial ANOVA for the AID β-gal expression did not reveal any main effects or an interaction (see Figure 3c). In contrast, β-gal expression in the AIV differed as a function of Ethanol Dose (F_2,51_ =3.598, *p*<0.05), with a significant decrease in β-gal expression relative to saline controls evident in animals conditioned with 2.5 g/kg ethanol, when collapsed across Adolescent Exposure condition (see Figure 3d). However, one-way follow-up ANOVAs conducted separately for each Adolescent Exposure did not reveal main effects of Ethanol Dose.

##### Nucleus Accumbens Core (NAcC), Basolateral Amygdala (BLA) and Central Amygdala (CeA)

The factorial ANOVAs of β-gal expression in the NAcC, BLA, and CeA did not reveal main effects of Exposure, Dose, or interactions of Exposure X Dose (see Figure 4 e, f, g).

**Figure 4.**
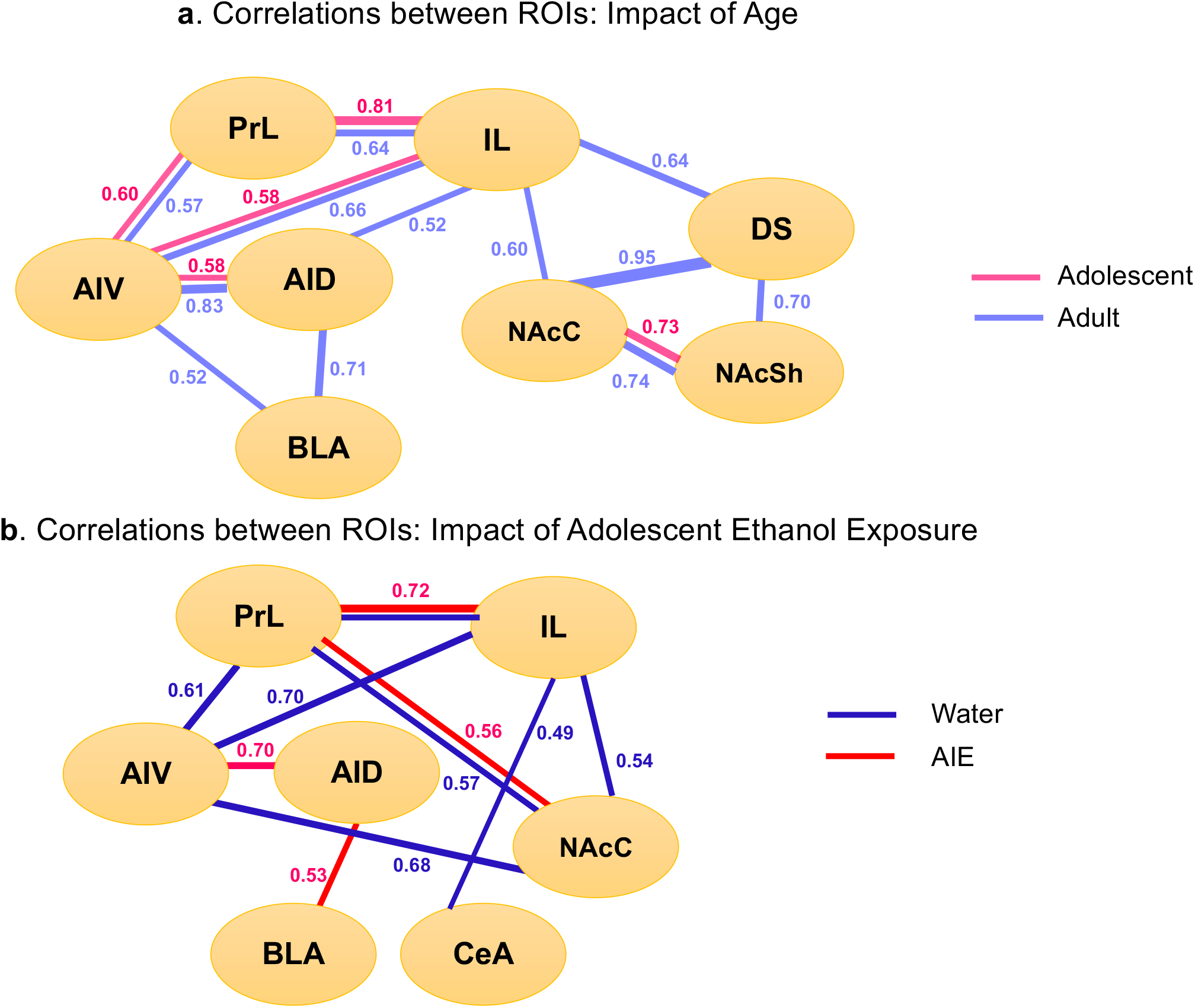
Correlation matrix as a measure of functional connectivity between brain regions in adolescents and adults (a) and water and AIE exposed adults (b). Only significant (p <0.05) correlations are presented, number represent r values.

##### Forward-Stepwise Multiple Regression: β-gal expression and Test Day intake

In AIE exposed animals, the forward stepwise multiple regression model revealed a significant positive relationship between β-gal expression in the AID and test day intake (adjusted R^2^=0.435, *p*=0.007), with greater β-gal expression in the AID related to greater intake of the CS on test day. There were no other significant predictors of test intake in the AIE group. In water exposed subjects, there were significant positive relationships between test day intake and β-gal expression in the AIV (adjusted R^2^=0.639, *p*=0.022) and AID (adjusted R^2^=0.639, *p*=0.039). All other ROIs did not differ from the model by the stepwise procedure.

##### Patterns of neuronal activation: Correlation matrix

As in Experiment 1, Pearson bivariate correlations were conducted using pairwise comparisons of β-gal+ cell counts in each ROI (see Figure 4b). In water and AIE exposed animals, significant positive correlations (all *p* < 0.05) were evident between the PrL and IL as well as the PrL and NAcC. In the water-exposed condition, significant positive correlations were also evident between the PrL and AIV, IL and AIV, AIV and NAcC, IL and NAcC, as well as the IL and CeA. In AIE animals, significant positive correlation were evident between the AID and AIV as well as the AID and BLA.

## Discussion

The results of the present experiments demonstrate attenuated sensitivity to ethanol-induced CTA in adolescent males relative to their adult counterparts as well as adult male rats following AIE exposure in early adolescence relative to water exposed control males. While adults in Experiment 1 and water-exposed controls in Experiment 2 demonstrated CTA following conditioning with both doses of ethanol, adolescents and AIE-exposed adults developed CTA only following the highest dose. The observed age-related difference is in agreement with previous studies that have shown attenuated sensitivity to ethanol-induced CTA in adolescent male rats (Anderson et al., 2014; Anderson et al., 2010; Holstein et al., 2011; Moore et al., 2013; Morales et al., 2014). These age-related differences in sensitivity to the aversive effects of ethanol are likely attributable to several cellular processes underlying CTA learning and/or expression rather than other physiological consequences of acute ethanol challenge such as hypothermia, sedation, or blood and brain ethanol (Schramm-Sapyta et al., 2010). Similarly, the results of Experiment 2 agree with previous studies that have shown attenuated CTA to ethanol following AIE exposure in adult male mice and rats (Diaz-Granados & Graham, 2007; Saalfield & Spear, 2015). Importantly, attenuated sensitivity to ethanol-induced CTA is evident following different modes of adolescent ethanol exposure: intragastric (Saalfield & Spear, 2015), intraperitoneal (Alaux-Cantin et al., 2013), and vapor (Diaz-Granados & Graham, 2007) exposures to ethanol during adolescence result in decreased sensitivity to the aversive effects of ethanol. While demonstrating similar insensitivity to ethanol-induced CTA, adolescents and AIE exposed adults had different patterns of neuronal activation and functional connectivity.

Distinct age-related differences in β-gal expression were evident in Experiment 1, with adolescents having a significantly greater number of β-gal+ cells than adults in all ROIs except for the BLA. This may reflect a greater number of excessive synaptic connections and neurons in young adolescents relative to adults prior to synaptic pruning, a hallmark of adolescent development (Spear, 2013). Both ages, however, showed very limited CTA-associated changes in neuronal activation across ROIs when re-exposed to the CS. In adolescents, the CeA was the only region affected by previous conditioning with the ethanol US: β-gal expression was significantly decreased in adolescents conditioned with ethanol. Given that the CeA is a hub for anxiety and alcohol circuits (Gilpin et al., 2015), the observed decreases in β-gal expression suggest that adolescents, but not their adult counterparts, might experience anxiolytic effects of ethanol during conditioning which resulted in decreased anxiety and attenuated activation of the CeA on test day. In contrast to adolescents, non-manipulated adults showed increases in β-gal expression in the PrL, IL, and AIV when conditioned with a 1.5 g/kg ethanol dose. This finding is in agreement with previously reported relationships between immediate early gene expression and aversion strength: increased c-fos expression in the insular cortex was evident in animals with moderate, but not strong CTA (Hadamitzky et al., 2015). Furthermore, c-Fos expression in the PrL and IL has been found following complete extinction of CTA (Mickley et al., 2005), with protein synthesis inhibition in the mPFC also impairing CTA extinction (Akirav et al., 2006). Therefore, increased β-gal expression in the insular cortex, PrL, and IL of adult males that were conditioned with the 1.5 g/kg ethanol dose may reflect, to some extent, extinction of CTA, especially given that animals were allowed to ingest NaCl for 60 minutes.

Age-related differences observed in Experiment 1 differ from those reported in the only published study that assessed c-Fos activation patterns associated with ethanol-induced CTA in adolescents and adults (Saalfield & Spear, 2019). Saalfield and Spear (2019) examined c-Fos induction in response to re-exposure to the US (i.e., challenge dose of EtOH) following CS-US pairing and found age differences in baseline and ethanol-induced c-Fos expression. Specifically, under baseline conditions, adolescents had significantly less c-Fos positive neurons than adults in the PrL but more than adults in the bed nucleus of the stria terminalis (BNST). Adolescent-typical ethanol-induced c-Fos increases were evident in the NAcC and NAcSh, whereas ethanol elicited c-Fos expression in the BNST regardless of age. The discrepancies between the studies may be related to the fact that we exposed animals to the CS, whereas adolescents and adults were re-exposed to the US in the Saalfield and Spear study. Indeed, exposure to the CS (Glover et al., 2016; Navarro et al., 2000; Yasoshima et al., 2006), US (Sakai & Yamamoto, 1997) or initial CS-US pairing using LiCl (Ferreira et al., 2006) result in different patterns of neuronal activation assessed via c-Fos expression. In the present work, the reason for measuring β-gal (proxy of c-Fos) expression in response to the CS as opposed to the US (ethanol) after CS-US pairing was to avoid/reduce β-gal induction not specifically associated with the aversive effects of ethanol. Ethanol produces multifaceted effects by affecting brain circuits involved in reward, motivation, and stress (Rhinehart et al., 2020), therefore it may be difficult to discern whether β-gal expression in response to the US is reflective of aversive or some other property of EtOH, such as level of intoxication (Bachtell et al., 1999; Sharpe et al., 2005). Moreover, the present study used cFos-LacZ transgenic rats on a Sprague-Dawley background as opposed to wildtype Sprague Dawley rats to investigate neuronal activation via β-gal expression. β-gal is induced only in strongly activated Fos+ neurons and not in surrounding nonactivated (Fos-) or weakly activated neurons (Cruz et al., 2014).

In Experiment 2, the IL was the only brain region in which a difference in β-gal expression between AIE and water-exposed subjects occurred, with a significantly greater number of β-gal+ cells in the AIE exposed rats relative to their water exposed counterparts when collapsed across EtOH doses. It is likely that greater β-gal expression in the IL of AIE-exposed males is associated with elevated levels of anxiety in these animals, given that AIE exposure results in enhanced anxiety-like behavioral alterations (reviewed in Towner & Varlinskaya, 2020) and the imbalance between excitation and inhibition toward excitation (Healey et al., 2021; Swartzwelder et al., 2017), as activation of pyramidal neurons in the IL enhances anxiety responding by shifting the balance to excitation (Berg et al., 2019), Significant effects of ethanol were evident in the IL and AIV, with ethanol decreasing β-gal expression when collapsed across adolescent condition. These findings were rather unexpected, since adult non-manipulated subjects demonstrated greater β-gal expression in these ROIs following conditioning with the 1.5 g/kg ethanol dose.

Age differences (Experiment 1) and AIE effects (Experiment 2) became more apparent when the correlation analysis was used. Although this analysis revealed a circuit involving connections between the PFC (PrL, IL) and agranular insula (AI) as well as between subregions of the PFC (i.e., PrL and IL) and AI (i.e., AID and AIV) in both age groups, adolescents had substantially less functional connections between the ROIs than adults (see Figure 4a). In adults, there were two additional circuits of functional connections that were not evident in adolescent rats. The first circuit included functional connections between the IL, DS and NAc, whereas the second one included the AID, AIV and BLA. Together, these findings suggest rather limited connected involvement of cortical with subcortical brain regions in responsiveness to the CS in adolescent males relative to their adult counterparts, although cortical ROIs appear to be active regardless of age. The correlation analysis also revealed substantial differences in functional connectivity between adult AIE-exposed males and their water-exposed counterparts. As in Experiment 1, water-exposed controls showed significant positive correlations between the AIV and PFC (i.e., PrL, IL), whereas these correlations did not reach significance in AIE animals. In contrast, functional connectivity between the two subdivisions of the agranular insula (AIV and AID) was evident in AIE-expose rats, but not in their water-exposed counterparts. Furthermore, the correlation analysis revealed a circuit involving connections between the IL, AIV, and NAcC as well as a functional connection between the IL and CeA only in water-exposed controls, with the AID and BLA functional connection evident only in AIE males. These findings suggest rather limited involvement of cortical brain regions in responsiveness to the CS following AIE.

Together, the results of Experiments 1 and 2 further support the roles of the PFC, agranular IC, nucleus accumbens and amygdala in responsiveness to the CS in EtOH-induced CTA. Furthermore, although adolescents and AIE exposed adults demonstrated different patterns of β-gal expression and functional connectivity, the stepwise multiple regression revealed similar positive relationships between β-gal+ neurons in the AID and CS intake on test day in these animals as well as non-manipulated and water exposed adults, confirming an important role of the insular cortex in responding to a tastant (Stehberg et al., 2011). These findings are also in agreement with previous studies that have demonstrated an important role of the PFC in both the acquisition and extinction of CTA (Gonzalez et al., 2015; Hernádi et al., 2000; Mickley et al., 2005). The results of the correlation analysis confirm previously shown involvement of the amygdala and nucleus accumbens in CTA (Yasoshima et al., 2006). According to Yamamoto (2007), the CeA facilitates detection of the CS, whereas the BLA is implicated in the hedonic shift from positive to negative. Furthermore, synchronized activity of the PrL, agranular IC, and BLA has been shown after pairing of a tastant and LiCl (Uematsu et al., 2015). In addition, functional connections between the insular cortex and BLA regulates CTA memory allocation in the insular cortex (Abe et al., 2020), with communication between cortico-amygdalar structures being important for CTA acquisition (Huang et al., 2020; Kayyal et al., 2019; Lavi et al., 2018). It has been shown that activation of neurons in the anterior insular cortex projecting to the BLA in adult male mice is necessary for forming memories that associate taste with negative valence (Kayyal et al., 2019). Increases of c-Fos have been observed in the insula, NAc (both shell and core), and BLA following pairing of saccharin and i.p. LiCl (Soto et al., 2017), suggesting functional connectivity between all these regions at least during CTA acquisition.

The main limitation of the present study is the use of only male subjects. Just a few studies have investigated the effect of sex on ethanol-induced CTA, with some evidence that sex differences may be affected by age and social context (Schramm-Sapyta et al., 2014; Vetter-O’Hagen et al., 2009) as well as adolescent ethanol exposure (Sherrill et al., 2011). Further studies are needed for assessment of possible age differences and effects of AIE on sensitivity to the aversive effects of ethanol in females. Another limitation of the current study is that subtypes of activated neurons cannot be identified with X-gal staining. Identification of the subtypes of cells activated during re-exposure to the CS can provide valuable information about mechanisms implicated in the aversive properties of ethanol.

In summary, our results confirm relative insensitivity of adolescent male rats to the aversive effects of ethanol. This relative insensitivity of adolescents may be associated with attenuated functional connectivity between cortical regions (prelimbic, infralimbic, insular) and the amygdala and nucleus accumbens core. Exposure of male rats to ethanol during adolescence results in insensitivity to the aversive effects of ethanol later in life, with attenuated functional connectivity between the PFC, IC, and NAcC being similarly involved in this insensitivity.

## Acknowledgements

The work presented in this manuscript was funded by NIH grants U01 AA019972 (Neurobiology of Adolescent Drinking in Adulthood Consortium - NADIA Project) and P50AA017823. Any opinions, findings, and conclusions or recommendations expressed in this material are those of the author(s) and do not necessarily reflect the views of the above stated funding agencies. The authors have no conflicts of interest to declare.

